# Opto-Katanin: An Optogenetic Tool for Localized Microtubule Disassembly

**DOI:** 10.1101/2021.12.22.473806

**Authors:** Joyce C. M. Meiring, Ilya Grigoriev, Wilco Nijenhuis, Lukas C. Kapitein, Anna Akhmanova

**Affiliations:** Cell Biology, Neurobiology and Biophysics, Department of Biology, Faculty of Science, Utrecht University, Utrecht, Netherlands; Center for Living Technologies, Eindhoven-Wageningen-Utrecht Alliance, the Netherlands

## Abstract

Microtubules are major cytoskeletal filaments that drive chromosome separation during cell division, serve as rails for intracellular transport and as a scaffold for organelle positioning. Experimental manipulation of microtubules is widely used in cell and developmental biology, but tools for precise subcellular spatiotemporal control of microtubule integrity are currently lacking. Here, we exploit the dependence of the mammalian microtubule-severing protein katanin on microtubule-targeting co-factors to generate a light-activated system for localized microtubule disassembly that we named opto-katanin. Targeted illumination with blue light induces rapid and localized opto-katanin recruitment and local microtubule depolymerization, which is quickly reversible after stopping light-induced activation. Opto-katanin can be employed to locally perturb microtubule-based transport and organelle morphology in dividing cells and differentiated neurons with high spatiotemporal precision. We show that different microtubule-associated proteins can be used to recruit opto-katanin to microtubules and induce severing, paving the way for spatiotemporally precise manipulation of specific microtubule subpopulations.

## Introduction

Microtubules have been implicated in nearly every step of cell physiology. They separate chromosomes during cell division, serve as tracks for long-range intracellular transport and organelle positioning, control cell and organelle morphology, and regulate adhesion turnover in migrating cells ^1 2 3^. Given the broad significance of microtubules, tools for their discrete manipulation are highly sought after to study the role of microtubules in specific processes while minimizing global cell perturbation. Recent development of light-activatable microtubule-targeting drugs enabled inhibition of microtubule polymerization in single cells ^4-6^. However, subcellular microtubule disruption with such drugs was thus far only achievable in long neuronal protrusions ^5 6^, because these compounds revert to an inactive state slowly, if at all, and diffuse rapidly out of the light-targeted area ^7^. Laser ablation has also been used to perform localized microtubule cutting within cells ^8, 9^. It is an effective tool for targeted ablation of diverse cellular structures, notably the destruction of centrosomes ^9^. However, this technique does not discriminate between microtubules and other cellular components and can therefore cause undesired side-effects. Optogenetic approaches have been very successful in exerting both specific and targeted subcellular effects, such as steering of migrating cells by controlling protein composition at growing microtubule ends using photo-inactivated End Binding protein 1 (EB1) ^10^.

Here, we set out to achieve potent and specific localized microtubule disruption by developing an optogenetic tool for microtubule severing. Optogenetic tools are built using engineered proteins that undergo light-induced conformational changes leading to altered protein activity, dimerization, multimerization or dissociation. This activity is facilitated by a photosensory module, which changes conformation upon illumination with light of specific wavelength. Widely used photosensory modules with the smallest size are the Light-Oxygen-Voltage (LOV) family of protein domains. The improved Light Induced Dimerizer (iLID) system consists of an optimized LOV2 domain derived from *Avena sativa* conjugated with the bacterially derived SspB-binding peptide SsrA ^11^. SsrA is sterically blocked when LOV2 is in the dark state but becomes uncaged upon blue-light activation of LOV2 and available to bind SspB. Several different SspB modules with different binding affinities have been engineered for the iLID-SspB heterodimerization system ^11, 12^, making it easy to apply and optimize for different applications. The iLID-SspB system also offers a larger dynamic range than older systems, more than 50-fold increase in affinity upon activation, and does not oligomerize thus avoiding aggregate-induced perturbations of cell functions ^11^.

Previously, an optogenetic microtubule-disrupting tool was developed in the form of a light recruitable microtubule catastrophe promoter, kinesin-13 ^13^. However, inducible recruitment of kinesin-13 was not sufficient to locally deplete all microtubules, likely because kinesin-13 triggers disassembly of growing microtubule ends but is less efficient in depolymerizing stable microtubule lattices ^14^. Since we wanted to develop a tool that could also disassemble microtubule lattices, we opted to use a microtubule severing enzyme instead. We combined the iLID-SspB system with the microtubule severing enzyme katanin. Katanin consists of p60 and p80 subunits, where p60 is the enzymatically active subunit that forms a hexamer which interacts with β-tubulin tails to extrude tubulin dimers from the microtubule lattice ^15-18^. Previous reports showed that mammalian p60 has low microtubule binding affinity on its own and requires co-factors like katanin p80, CAMSAP3 and ASPM for its recruitment to microtubules and microtubule-severing activity ^19-21^. Since p60 severing activity depends on microtubule recruitment, we considered this enzyme to be an optimal candidate for an optogenetic tool. We tested an assortment of microtubule-binding proteins and domains to arrive at an optimal microtubule binding anchor, an EB3 microtubule-binding domain with a VVD_fast_ based reversible blue light-sensitive homodimerization domain ^22^. In utilizing two separate blue light-sensitive systems, VVD_fast_ to control recruitment of the anchor to microtubules, and iLID-SspB to control the recruitment of katanin to microtubules, we were able to minimize dark state activity and off-target binding in order to limit the impact on normal microtubule architecture. Our system, which we named opto-katanin, was able to achieve localized and reversible microtubule disassembly within 1.5 min in cultured cancer cells and 10 min in neuronal protrusions. We further demonstrated that opto-katanin-mediated microtubule severing could be combined with imaging of organelles to observe the impact of local microtubule disassembly on transport and organelle morphology.

## Results

### Localized and reversible microtubule disassembly by light-controlled katanin recruitment

In order to make a system that could recruit katanin to microtubules in a light-dependent manner, we designed a microtubule anchor incorporating iLID and a p60 construct linked to SspB_micro_. The microtubule binding domain of EB3 (EB3_N_) was used as a microtubule anchor. Artificial EB dimer constructs have been reported to bind the microtubule lattice with considerably higher affinity than artificial EB monomers ^23, 24^. Therefore we incorporated a blue light sensitive Vivid (VVD) optogenetic homodimerization domain containing Ile52Cys mutation for improved homodimer stabilization ^25^ and Ile74Val and Ile85Val mutations to increase the rate of reversibility ^22, 26, 27^, also known as VVD_fast_ ^22^. With a two-step activation consisting of dimerization of VVD and unfolding of iLID being necessary for efficient activation of the construct we hoped to reduce dark state activity. This also aimed to reduce competition of the anchor construct with endogenously expressed microtubule associated proteins. The EB3_N_ microtubule anchor and p60 construct were labelled with mClover3 or HaloTag and mCherry, respectively, so that their expression levels and localization could be assessed (Fig. 1a). The HaloTag anchor was used so that anchor localization could be visualized with a far red HaloTag dye without activating iLID-SspB and VVD_fast_. Flexible GGGGS linkers were inserted between protein domains to support proper protein folding and function (Fig. 1a). We expected that blue light application would trigger homodimerization of EB3_N_-VVD-mClover3-iLID or EB3_N_-VVD-Halo-iLID, which would enhance microtubule binding affinity, and simultaneously unfold iLID allowing binding of SspB-mCh-p60, recruiting it to microtubules and thus prompting their disassembly (Fig. 1b).

**Fig. 1:**
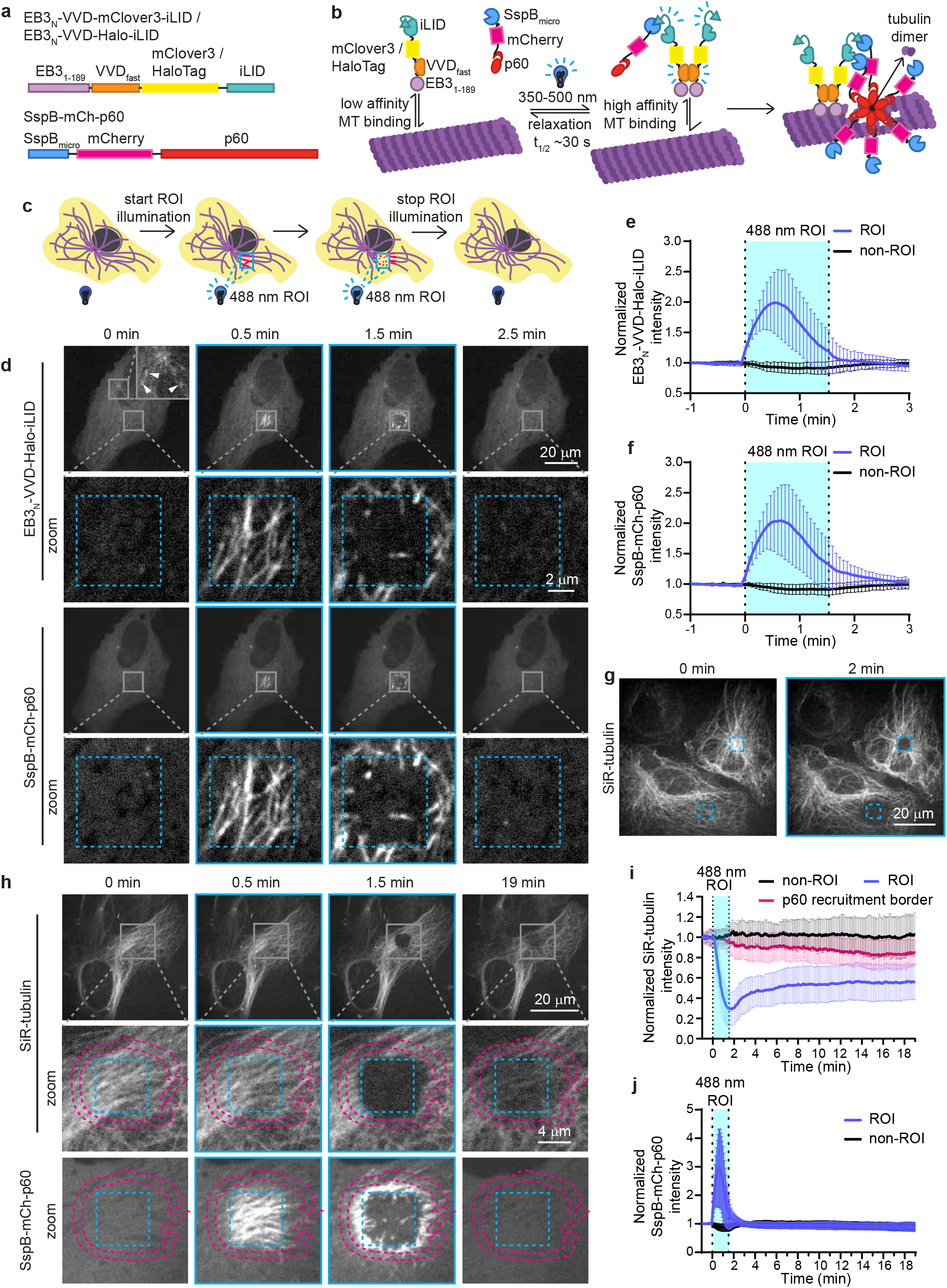
Converting katanin into an optogenetic tool for localized and reversible microtubule disassembly. **a**, Overview of optogenetic constructs; protein domains are separated by flexible GGGGS linkers. **b**, A scheme showing the mechanism of microtubule disassembly by opto-katanin. In the dark state, EB3_N_-VVD-mClover3-iLID or EB3_N_-VVD-Halo-iLID constructs bind microtubules with low affinity and iLID does not bind SspB. Upon blue light activation, VVD_fast_ homodimerizes enabling high affinity microtubule binding of EB3 microtubule binding domain. Blue light also causes iLID to change conformation, revealing a SsrA domain that engages with SspB thus recruiting SspB-mCh-p60 to microtubules and promoting microtubule disassembly by p60. **c**, Scheme of experimental set up: cells co-transfected with optogenetic constructs are locally pulsed with blue light to induce localized construct recruitment and microtubule disassembly. Blue light pulsing is then stopped to observe the off rate of optogenetic constructs and local microtubule recovery. **d**-**j**, U2OS cells co-transfected with SspB-mCh-p60 and either (**d**-**f**) EB3_N_-VVD-Halo-iLID or (**g**-**j**) EB3_N_-VVD-mClover3-iLID, with (**g**) 200 nM, (**h**-**j**) 100 nM or (**d**-**f**) without SiR-tubulin labelling were imaged on a spinning disc microscope. **d**-**f, h**-**j**, Cells are imaged for 0.5 min before locally pulsing with blue light inside ROI (marked by blue dashed box) for 1.5 min and subsequently imaged without blue light pulsing. **d, h**, Representative stills at the indicated time points before the start of blue light illumination. **g**, Stills of cells before and after 2 min of blue light pulsing within 2 ROI (marked by blue dashed boxes). **e, f, i, j**, Quantification of fluorescence intensity inside blue light pulsed areas (ROI) and comparable sites not pulsed with blue light (non-ROI) in the same cell for the indicated channels normalized to the average of the first 5 time points. n = 17-18 cells across 3 independent experiments, graphs show mean ± SD.

Prior to blue light illumination, EB3_N_-VVD-Halo-iLID was slightly enriched at microtubule plus ends. This fits with previous observations that the monomeric microtubule binding domain of EB3 still shows some affinity for growing microtubule plus ends ^24^ (white arrows, Fig. 1d and Supplementary Video 1). SspB-mCh-p60 was observed to be enriched at the centrosome but was mostly cytoplasmic, which is consistent with prior localization studies of p60 (Fig. 1d and Supplementary Video 1). Upon localized blue light pulsing in a region of interest (ROI), both EB3_N_-VVD-Halo-iLID and SspB-mCh-p60 were recruited to microtubules in the ROI, reaching peak enrichment approximately 30 s after commencing blue light pulsing, before intensity dropped back due to microtubule severing (Fig. 1d, e, f and Supplementary Video 1). After microtubules were disassembled inside the ROI, EB3_N_-VVD-Halo-iLID and SspB-mCh-p60 became enriched on microtubules bordering the ROI, due to diffusion of activated constructs in the absence of microtubules to bind (Fig. 1d). If blue light illumination was continued, microtubules bordering the ROI were also disassembled, and to prevent this, illumination was stopped after 1.5 min, just after microtubules inside the ROI were disassembled. Constructs redispersed within 1 min of stopping blue light application, indicating fast reversibility (Fig. 1d, e, f and Supplementary Video 1). In addition to SspB-mCh-p60, a construct containing the higher affinity SspB_nano_ instead of SspB_micro_ was also tested, but showed undesirable dark state activity and microtubule localization prior to activation.

In order to confirm that microtubules were locally disassembled, opto-katanin-mediated microtubule severing was performed in cells labelled with SiR-tubulin. Indeed, opto-katanin severing could clear an area of SiR-tubulin labelled microtubules, both at the cell periphery and near the centrosome (Fig. 1g, Supplementary Video 2). To test microtubule recovery after severing, we switched to the lowest usable concentration of SiR-tubulin to label microtubules while minimizing the microtubule stabilizing activity of the SiR-tubulin probe. As expected, the loss of SspB-mCh-p60 from the ROI after activation correlated with a steep decrease in SiR-tubulin intensity indicative of microtubule severing (Fig. 1h-j, Supplementary Video 3). Reduction in SiR-tubulin appeared extremely localized, suggesting that the area of microtubule damage could be contained well (Fig. 1h). To confirm this, we measured SiR-tubulin intensity just outside the border of where p60 was recruited to microtubules (p60 recruitment border, pink dotted line, Fig. 1h) and found that it reduced only slightly after microtubule severing (Fig. 1i). These data indicate that opto-katanin-mediated severing does not result in significant microtubule disassembly outside the illuminated area, consistent with *in vitro* work showing that katanin-mediated severing leads to spontaneous incorporation of GTP-tubulin and increases microtubule rescues ^28^. This contrasts with laser ablation-mediated microtubule severing, which induces extensive microtubule depolymerization ^8^. The recovery of SiR-tubulin signal inside the ROI was observed 6 min after stopping light activation, and this signal reached a plateau at approximately 33% of the original intensity (Fig. 1i), possibly because microtubules were not sufficiently dynamic to repopulate the area on this timescale. We cannot rule out that despite efforts to minimize SiR-tubulin-induced microtubule stabilization, there is still some stabilizing effect that would be expected to delay recovery ^29^.

### Opto-katanin microtubule disassembly with different microtubule anchors

We next used several different MAPs to test whether other microtubule anchors could be employed and what properties made a good anchor for the opto-katanin tool (Table 1). Various microtubule anchors were made and tested in live cells co-transfected with SspB-mCh-p60, assessed in live cell imaging experiments and scored for microtubule binding, bundling and severing (Table 1 and Extended Data Fig. S1). From the anchors tested it was apparent that microtubule bundling activity, such as observed for MAP7, Tau and MAP2C, was generally predictive of poor microtubule severing activity, despite these anchors being able to recruit katanin (Table 1 and Extended Data Fig. S1). The observation that the shorter MAP7 constructs missing the C-terminal domain did not make microtubule bundles and do perform well as microtubule severing anchors (Table 1) further supports the notion that microtubule bundling activity inhibits opto-katanin mediated severing. As to be expected, the absence of microtubule binding was also a good predictor of poor severing activity, since such anchors cannot effectively recruit SspB-mCh-p60 to microtubules (Table 1 and Extended Data Fig. S1). We found that the addition of mClover3-iLID could reduce microtubule binding affinity and in some cases resulted in abrogation of microtubule binding (Table 1). There were several instances of constructs using previously identified microtubule binding domains showing poorer microtubule localization compared to the full-length constructs, for example, MAP7D3, Tau0N4R^30^ and MAP2C^31^ (Table 1). This implies that other protein domains contribute to microtubule binding, and indeed, previous work has suggested that MAP7D3 also contains a microtubule binding region in its C-terminus^32, 33^. This result also agrees with findings that the 4R microtubule binding domain of Tau alone was previously found to have a 25-fold reduction in microtubule binding affinity compared to full-length Tau ^30^.

**Table 1:** Performance of anchor constructs in microtubule binding, bundling and severing. Severing performance was assessed in the context of co-transfection with SspB-mCh-p60 and live cell imaging with blue light illumination. ++ high, + moderate, +/- poor, - not observed. Constructs with an * are shown in Fig. 2. See Fig. S1 for details on grading metrics.

| Anchor Construct | MT binding | MT bundling | MT severing |
| --- | --- | --- | --- |
| EB3<sub>N</sub> (1-189)-VVD<sub>fast</sub>-mClover3-iLID * | + dark/++ lit | - | ++ |
| EB3<sub>N</sub> (1-189)-VVD<sub>fast</sub>-Halo-iLID * | + dark/++ lit | - | ++ |
| iLID-mClover3-DCX * | ++ | - | ++ |
| iLID-mClover3-MAP7 | ++ | ++ | +/- |
| iLID-mClover3-MAP7<sub>N</sub>(1-301) * | ++ | - | ++ |
| iLID-mClover3-MAP7<sub>MTBD</sub>(59-170) | ++ | - | ++ |
| iLID-mClover3-MAP7D3 * | ++ | - | ++ |
| iLID-mClover3-MAP7D3<sub>N</sub>(1-309) | + | - | ++ |
| iLID-mClover3-MAP7D3<sub>MTBD</sub>(55-174) | + | - | ++ |
| CLIP115<sub>N</sub>(1-309)-mClover3-iLID * | + | - | ++ |
| iLID-mClover3-Tau0N4R | ++ | ++ | +/- |
| iLID-mClover3-miniTau0N4R(140-342) | ++ | + | ++ |
| iLID-mClover3-Tau0N4R<sub>MTBD</sub>(186-310) | - | - | - |
| iLID-mClover3-Tau0N3R | ++ | ++ | +/- |
| iLID-mClover3-MAP2C | ++ | ++ | - |
| iLID-mClover3-MAP2C<sub>MTBD</sub>(300-393) | - | - | - |

We next focused in more detail on the anchors that did trigger MT light-induced disassembly by SspB-mCh-p60. These included proteins with different microtubule-binding sites: EB3_N_ ^34^ and Doublecortin (DCX) ^35^ that bind between protofilaments, MAP7_N_ ^36^ and MAP7D3, which straddle the inter-dimer and the intra-dimer tubulin interfaces, and the CAP-Gly domain containing CLIP115_N_, which binds the tail of α-tubulin ^37^ (Fig. 2a, b). Since excessive expression of constructs resulted in dark state activation of microtubule cutting, a maximum threshold for microtubule anchor and SspB-mCh-p60 expression was imposed. All five anchor constructs combined with SspB-mCh-p60 could induce microtubule depolymerization with whole cell blue light activation (LIT), visualized using α-tubulin antibody staining, which is much weaker in cells with disassembled microtubules (Fig. 2c, d). By using individual construct transfections and immunostaining, we confirmed that anchor constructs alone did not induce microtubule disassembly or reduce α-tubulin staining by blocking the antibody epitope (Extended Data Fig. S2). Certain anchors combined with SspB-mCh-p60 had a higher dark state activity than others; in particular, the CLIP115_N_ and the MAP7D3 anchors showed a large decrease in α-tubulin staining without light activation (Fig. 2d). EB3_N_ and MAP7_N_ anchors showed the lowest dark state activation (Fig. 2d). We confirmed that anchor and SspB-mCh-p60 expression was comparable for all conditions (Fig. 2e, f), therefore differences in microtubule density observed between DARK and LIT state and between different anchors could be attributed to optogenetic construct activation and differences in efficacy of the different anchors, respectively. Altogether, our data suggest that a good microtubule anchor for opto-katanin should bind individual microtubules well but not bundle them, and that a direct interaction with tubulin tails, like in CLIP115_N_, may be disadvantageous in this system. Since the EB3_N_ anchor exhibited the least dark state activity and the greatest difference between DARK and LIT conditions, this construct was used for subsequent experiments.

**Fig. 2:**
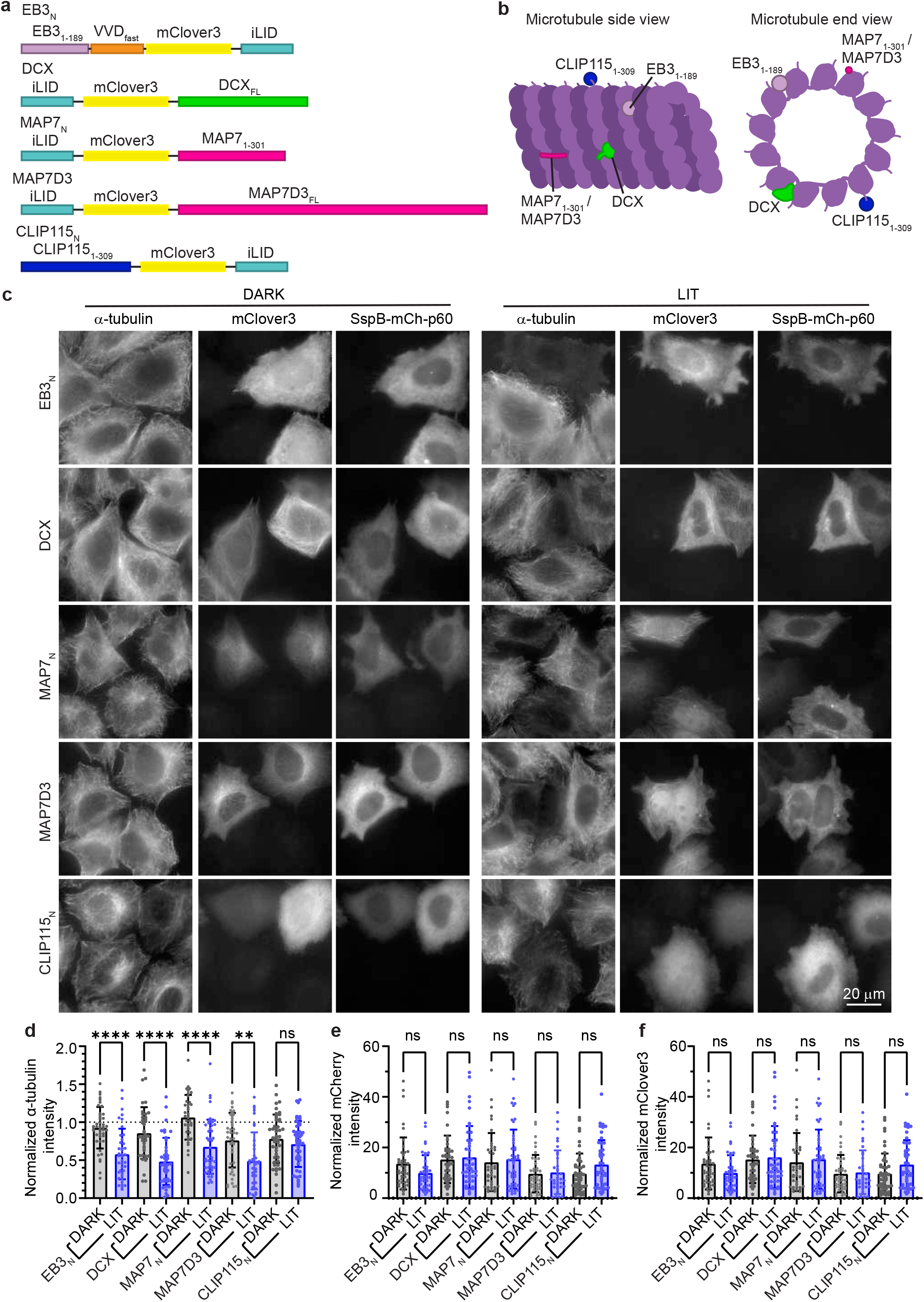
Different microtubule binding domains can support opto-katanin-mediated microtubule disassembly. **a**, Overview of optogenetic microtubule anchor constructs. **b**, Scheme showing the position of microtubule binding sites of EB3_N_, DCX, MAP7_N_/ MAP7D3 and CLIP115_N_ on the microtubule. **c**-**f**, HeLa cells were co-transfected with SspB-mCh-p60 and microtubule anchor constructs shown in (**a, b**) and either fixed in the dark (DARK) or fixed after a 16 minute incubation under blue light (LIT), prior to immunostaining for α-tubulin and imaging on a widefield microscope. **c**, Representative images. **d**-**f**, Quantification of normalized fluorescence intensity of (**d**) α-tubulin immunostaining, (**e**) mCherry intensity to indicate SspB-mCh-p60 expression or (**f**) mClover3 intensity to indicate microtubule anchor construct expression. **d**-**f**, n = 40-47 cells across 3 independent experiments, plots indicate mean ± SD with individual cell measurements shown as dot points.

### Localized opto-katanin microtubule severing to block intracellular transport

Having generated an effective tool for severing microtubules, we next wanted to test its application for locally disrupting microtubule functions, such as their ability to serve as tracks for transport. Among all cell types, neurons have the most elaborate and specialized transport network, which consists of a large population of stabilized microtubules bearing numerous post-translational modifications and decorated with a great diversity of MAPs. Given that a highly stable microtubule network may be difficult to disassemble, we first set out to test whether opto-katanin could sever microtubules in protrusions of mature day in vitro 9 (DIV9) neurons. We found that microtubules could be severed in neurites within approximately 10 min, as observed by SiR-tubulin labelling and the rise and fall of SspB-mCh-p60 intensity (Fig. 3a, b, c and Supplementary Video 4). Severing of acetylated and tyrosinated microtubule populations in both the neurites and cell body of fully illuminated neurons was also confirmed by antibody staining (Fig. 3d).

**Fig. 3:**
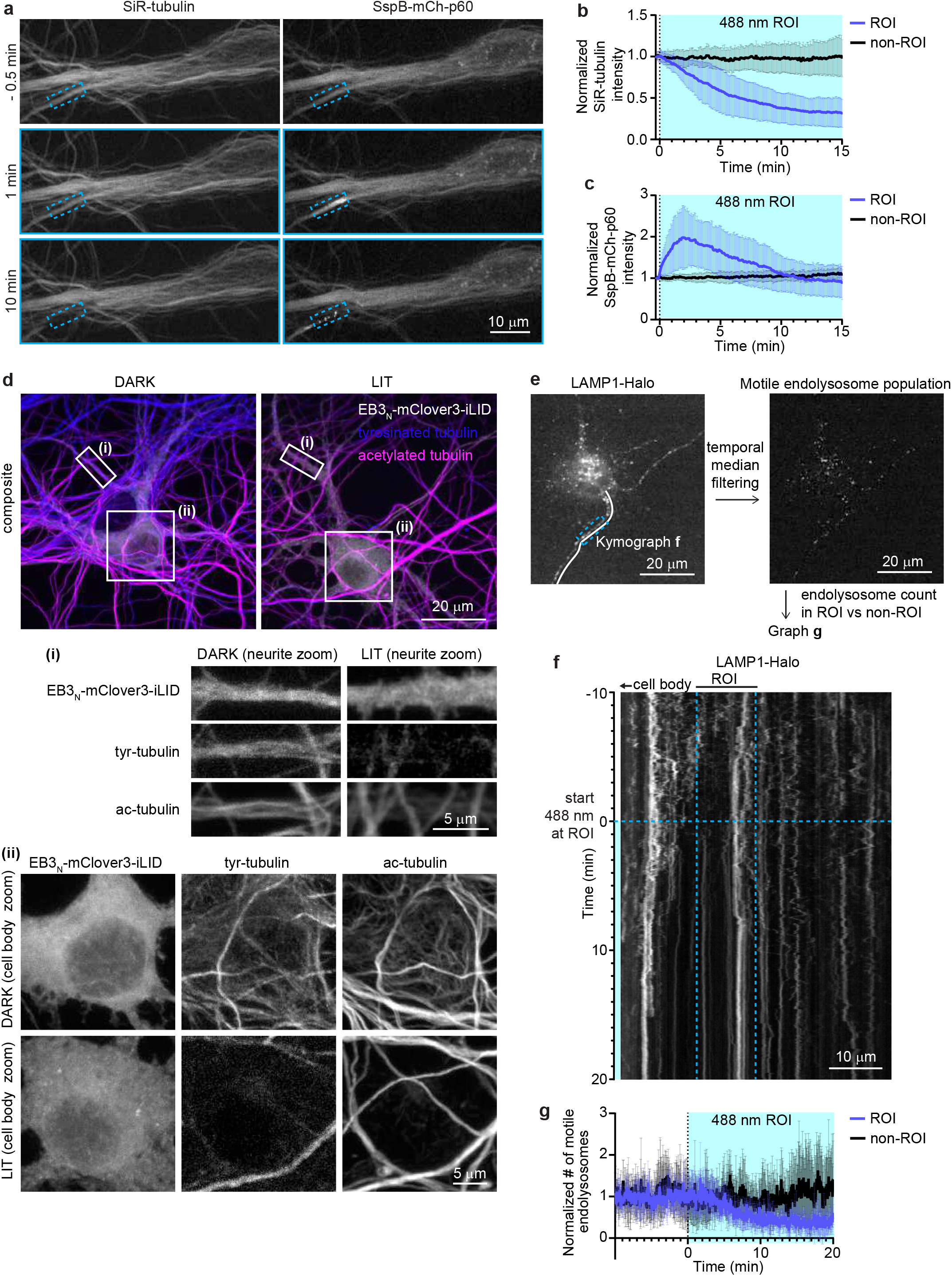
Localized optogenetic microtubule severing in neurons to block microtubule-based transport. **a**-**c**, DIV9 neuron co-transfected with EB3_N_-VVD-mClover3-iLID and SspB-mCh-p60 and stained with SiR-tubulin was pulsed locally with blue light at a ROI (marked by blue dashed box) from time = 0 min. **a**, Representative stills 0.5 min before (−0.5 min), 1 min after and 10 min after starting blue light pulsing. **b, c**, Quantification of normalized fluorescence intensity inside blue light pulsed region (ROI) and a comparable region in the same cell not illuminated with blue light (non-ROI) of (**b**) SiR-tubulin and (**c**) SspB-mCh-p60. Graphs indicate mean ± SD, n = 15-16 ROI from 8 neurons over 3 experiments. **d**, DIV9 neuron co-transfected with EB3_N_-VVD-mClover3-iLID and SspB-mCh-p60 either fixed in the dark (DARK) or illuminated with blue light for 16 minutes prior to fixation (LIT) and stained for tyrosinated tubulin (tyr-tubulin) and acetylated tubulin (ac-tubulin). White boxes show areas, (**i**) neurite and (**ii**) cell body, enlarged in zoom panels. **e**-**g**, DIV9 neuron co-transfected with EB3_N_-VVD-mClover3-iLID, SspB-mCh-p60 and LAMP1-Halo was pulsed locally with blue light (marked by blue dashed box/lines) from 0 min onward. **e**, LAMP1-Halo localization in the neuron with neurite shown in kymograph in (**f**) highlighted by a white line. Scheme of the analysis pipeline: temporal median filtering was applied to LAMP1-Halo movies to generate a movie showing only the motile endolysosome population, in which the motile endolysosomes were counted inside the blue light illuminated region (ROI) and in a comparable region not illuminated with blue light (non-ROI). **f**, Kymograph of LAMP1-Halo in neurite highlighted in (**e**). **g**, Quantification of motile endolysosomes as illustrated in panel (**e**); graph shows mean ± SD, n = 5 ROI from 4 neurons over 3 experiments.

In order to test the impact of microtubule severing on transport, we observed late endosome and lysosome (endolysosome) motility in DIV9 neurons using a LAMP1-HaloTag probe. As expected, the number of motile late endolysosomes dropped in the region where microtubules were locally disassembled, and in some cases, this could cause endolysosome traffic jams (Fig. 3e, f, g and Supplementary Video 5). Similarly, localized microtubule severing could also be used to block transport of exocytic Halo-Rab6A labelled vesicles in U2OS cells, where the vesicles paused near the microtubule cut site and then moved away along remaining microtubules (Supplementary Video 6).

### Targeted opto-katanin microtubule disassembly to perturb organelle morphology and dynamics

Aside from facilitating long-range transport, microtubules are also important regulators of the morphology of different organelles, including the Endoplasmic Reticulum (ER), Golgi and nucleus. We wanted to determine whether we could locally perturb organelle morphology using the opto-katanin tool. If the microtubule cytoskeleton is disassembled in cells using nocodazole, ER takes on a sheet morphology and loses tubular structures ^38, 39^. However, we observed that when microtubules were disassembled locally but maintained in the surrounding area (Fig. 4a), ER motility was locally inhibited and ER tubule density was reduced but the tubules did not transform into sheets (Fig. 4b, c, and Supplementary Video 7). Quantification demonstrated a localized drop in both motility and nodule density, which reflects ER tubule density, upon microtubule severing (Fig. 4d, e, f). Apparently, microtubule-based formation of ER tubules in the area surrounding the microtubule-depleted ROI is sufficient to maintain a tubular albeit less dense ER morphology within the microtubule-depleted region.

**Fig. 4:**
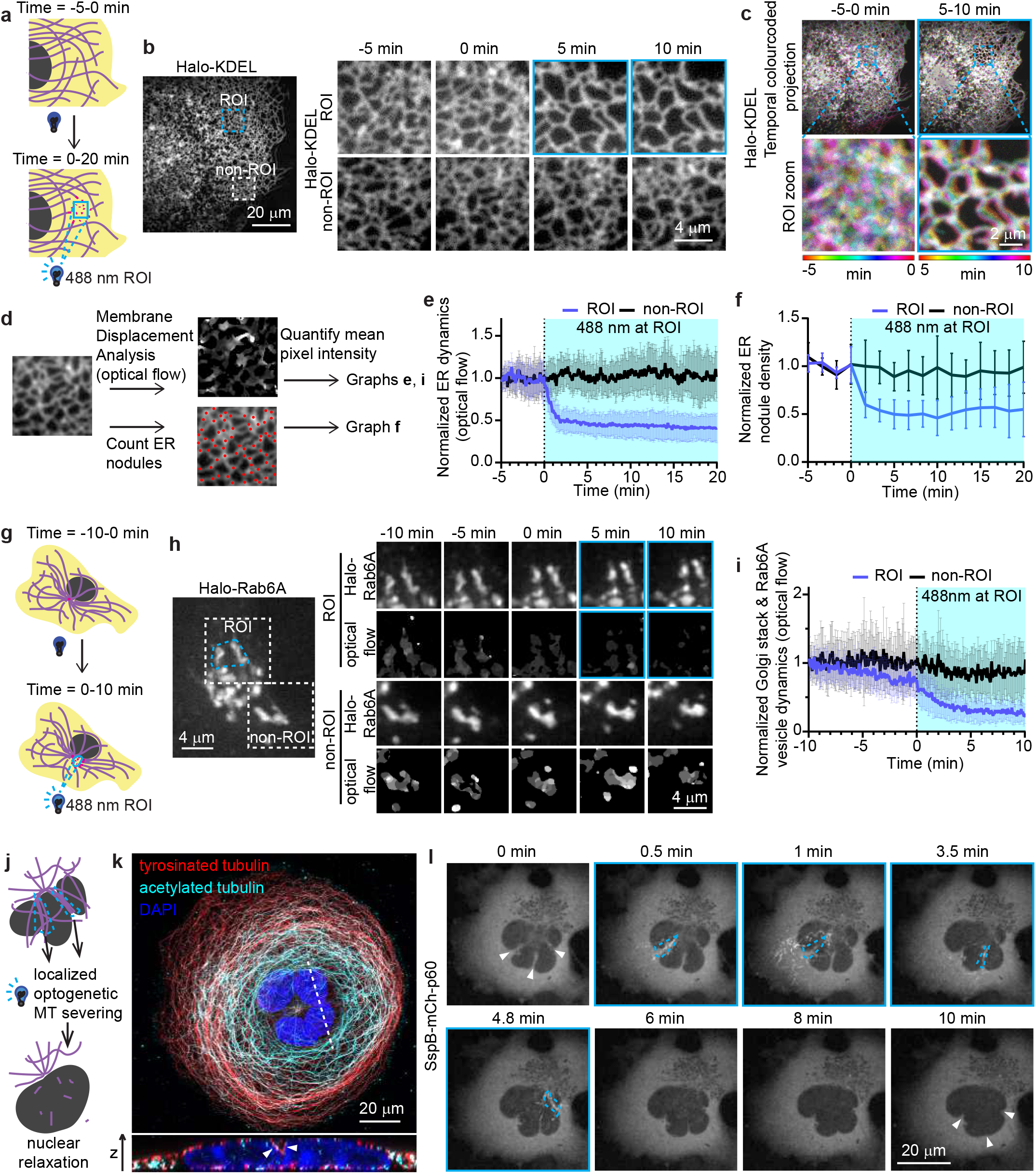
Localized optogenetic microtubule severing to perturb organelle morphology and dynamics. **a**-**f**, COS-7 co-expressing EB3_N_-VVD-mClover3-iLID, SspB-mCh-p60 and Halo-KDEL as an ER marker. **a**, Scheme of the experimental set up: microtubules were optogenetically severed in a central cytoplasmic region using localized pulses of blue light inside a ROI (marked by blue dashed box) from 0 min onward. **b**, Halo-KDEL in cell with zoomed in images of ROI and non-ROI control region at time points before microtubule severing (−5, 0 min) and after (5, 10 min). **c**, Z-projections of Halo-KDEL frames color coded for time before (−5-0 min) and after (5-10 min) local microtubule severing. **d**, Scheme of the analysis pipeline: membrane displacement was analyzed in ROI and non-ROI regions using optical flow Membrane Displacement Analysis that converts movement into pixel intensity, and then mean pixel intensity was quantified. ER nodules (marked by red dots) were also counted in ROI and non-ROI regions. **e, f**, Quantification of normalized ER dynamics (**e**) and normalized number of ER nodules (**f**), analyzed as illustrated in panel (**d**); graphs show mean ± SD, n = 16 cells across 4 independent experiments. **g**-**I**, U2OS co-expressing EB3_N_-VVD-mClover3-iLID, SspB-mCh-p60 and Halo-Rab6A to label Golgi and exocytic vesicles. **g**, Scheme of the experimental set up: ROI (marked by blue dashed box) was drawn around a subset of the Golgi stacks and pulsed with blue light from 0 min onward to locally sever microtubules. **h**, dynamics of Halo-Rab6A-labeled Golgi membranes; images show zoomed stills of Halo-Rab6A and the corresponding optical flow analysis (as illustrated in (**d**)) in ROI and non-ROI regions before (−10, −5, 0 min) and after (5, 10 min) local microtubule severing in ROI. **I**, Normalized dynamics of Halo-RAb6A-labeled membranes analyzed by optical flow; graph shows mean ± SD, n = 14 cells from 3 independent experiments. **j**, Scheme of the experimental set up: COS-7 with lobulated nuclei were pulsed with blue light sequentially within areas showing nuclear invaginations to sever microtubule bundles around the nucleus. **k**, Fixed COS-7 cell with a lobulated nucleus, stained for DAPI (dark blue), acetylated tubulin (cyan) and tyrosinated tubulin (red), showing microtubules inside nuclear invaginations (arrowheads). **l**, COS-7 cell co-expressing EB3_N_-VVD-mClover3-iLID and SspB-mCh-p60 with a lobulated nucleus was exposed to blue light in ROIs shown in dashed blue lines to sequentially sever microtubule bundles. After microtubule bundle severing, gradual nuclear relaxation was observed; arrows indicate locations of nuclear invaginations were present prior to microtubule severing.

Microtubules nucleated and anchored at the Golgi are used as tracks for vesicle transport ^2, 40^. While cells do not require microtubules for secretion ^41^, cooperative microtubule motor-based transport is necessary for long-distance translocation of secretory cargo ^42^. When microtubules were locally disassembled around a subset of (Rab6A labelled) Golgi stacks (Fig. 4g), the positioning of these stacks was not altered compared to the Golgi stacks in a region that was not illuminated with blue light (Fig. 4h, Supplementary Video 8). However, analysis of membrane dynamics (Fig. 4d) revealed a reduction of Golgi stack and Rab6A vesicle motility upon microtubule disassembly (Fig. 4h, i). This quantification indicates a decrease in lateral movement and transport, and does not rule out secretion of vesicles directly under the Golgi, as observed in kinesin motor knock out studies ^42^. Local microtubule disassembly also resulted in a local loss of Rab6A-positive Golgi tubules (arrows, Supplementary Video 8). This was consistent with previous reports that Golgi tubules align with microtubules and are extended as a result of kinesin-1-mediated pulling of Golgi membranes ^43^.

Microtubules can also participate in shaping the nucleus: in myeloid progenitors, microtubule bundles deforming the nucleus have been reported to influence chromatin organization and gene expression to drive differentiation ^3^. Microtubule-mediated nuclear constriction and formation of lobulated nuclei has also been observed in cancer cell lines and untransformed epithelial cells ^44, 45^. We observed lobulated nuclei in a subpopulation of COS-7 cells and tested whether it was possible to induce nuclear relaxation by cutting perinuclear microtubule bundles (Fig. 4j). Immunostaining revealed that microtubule bundles inside nuclear invaginations contained both tyrosinated and acetylated microtubules (Fig. 4k). We used opto-katanin to sequentially sever perinuclear microtubule bundles and observed slow sequential relaxation of nuclear invaginations, taking between 3-7 min after microtubule bundle severing (Fig. 4l, Supplementary Video 9). This suggests that, in contrast to the plastic deformation of nuclei in myeloid progenitor cells ^3^, microtubules were imposing constricting forces causing elastic deformation of COS-7 nuclei, which recovered a rounded shape after microtubule belts were cut. Furthermore, as observed in neurites, the opto-katanin tool could be applied to sever both acetylated and tyrosinated microtubules. Together, these data demonstrate that opto-katanin can be used to induce acute localized perturbations of organelle morphology and dynamics.

## Discussion

In this study, we have generated a potent tool for rapid, precise and reversible microtubule disassembly. Amongst different microtubule subtypes, we expected that stabilized populations such as acetylated microtubules and microtubule bundles would be the most difficult to sever, since they are the most resistant to mechanical stress and comprise the most long-lived population. In cultured cancer cells, stable and acetylated microtubules tend to be most abundant in the perinuclear area, and in mature neurons, they make up the vast majority of the microtubule population in neurites ^46^. However, we have demonstrated successful opto-katanin-mediated depletion of microtubule polymer in both the perinuclear area, neurites and perinuclear microtubule belts, evidenced by live SiR-tubulin labelling and immunolabelling of fixed cells. Profound microtubule disassembly in neurons is by itself is an important achievement, because complete microtubule depolymerization in neurons using conventional tools such as the tubulin-sequestering drug nocodazole is difficult or impossible ^47^. However, microtubule severing in neurites took significantly longer compared to severing in cultured U2OS or COS-7. Microtubules in mature neurons are strongly bundled and decorated with a variety of MAPs, properties that potentially inhibit katanin severing function. Increased stabilization may help to buffer the impact of local tubulin dimer extrusion from the lattice. Moreover, stabilizing factors may compete for microtubule binding with the optogenetic anchors and katanin itself, and microtubule bundling could potentially occlude access of the optogenetic severing constructs to microtubules. Indeed, from testing different microtubule anchors we noticed that microtubule bundling anchors such as Tau, MAP7_FL_ and MAP2C performed poorly despite their high microtubule binding affinity and ability to recruit katanin. In contrast, MAP7_N_, a MAP7 construct lacking the C-terminal part that supports microtubule bundling ^33^, served as a very efficient anchor for optogenetic katanin severing.

While we were preparing this study for publication, a similar inducible approach using another severing enzyme, spastin, was described ^48^. In this system, the authors used spastin mutations disrupting its binding to microtubules and a FRB/FKPB chemical heterodimerization or CRY2/CIBN light-inducible dimerization system to make the recruitment of mutant spastin to microtubules inducible ^48^. While most of this study was focused on the temporal control of spastin using a FRB/FKPB system, it also showed that local light-induced microtubule disassembly can be nicely achieved with the CRY2/CIBN based system though with the rate approximately 6 times slower than described here (∼9 min for spastin versus 1.5 min for opto-katanin). Further work will be needed for side-by-side comparison of the two systems, their dark state activity and off-target effects.

Future work may be done to make a severing tool specific for microtubule sub-populations. In designing the CAP-Gly domain CLIP115_N_ anchor, we attempted to make a tool to specifically disassemble tyrosinated microtubules. While unfortunately the CLIP115_N_ anchor had a high dark state activity and failed to show a good light induced microtubule severing response, additional MAPs or small antibodies with sensitivity for post translational modifications exist that could be explored as potential microtubule anchors ^49^. Indeed, FRB/FKPB-dependent spastin recruitment and microtubule severing could be induced by anchoring it with an antibody specific for tyrosinated tubulin, though it was not demonstrated that such a treatment leaves detyrosinated microtubules intact ^48^.

One consideration for use of opto-katanin is that if one is interested in using a fluorescent probe in combination with the tool, this probe is restricted to the far red or infrared channels. Fortunately, thanks to the development of HaloTag and SNAP-tag dyes, bright and stable options are available to image in far red. Infrared fluorescent proteins ^50^ would also be compatible for use with this tool. It is necessary to visualize SspB-mCh-p60 in the red channel because this makes it easy to identify transfected cells expressing optimal amounts of the construct to enable cutting without activating it. SspB-mCh-p60 also provides an important visual output indicating the location and size of the area of microtubule severing. Constructs where the mCherry tag in SspB-mCh-p60 were substituted for SNAP-tag or HaloTag have been tested, however both tags interfered with efficient microtubule severing.

We believe that opto-katanin is a strong asset for studying the functions of the microtubule cytoskeleton, since it is capable of performing precise, fast and reversible microtubule-specific manipulations that were previously impossible. We have demonstrated the application of this tool for acute cellular manipulations such as locally blocking microtubule motor-based transport, local perturbation of organelle morphology and dynamics and disruption of nuclear constriction. Potential research applications include the determination of the load-bearing contribution of microtubules in epithelial cell morphology ^51^, muscle contraction ^52^, nuclear positioning ^53^ and migration through soft substrates ^54^; studying microtubule orientation and assembly in neurites ^55^ and elucidating of the contribution of microtubules to various developmental morphogenetic processes.

## Online Methods

### Cell culture

HeLa, COS-7 and U2OS cells were cultured in Dulbecco’s Modified Eagle Medium (DMEM; Sigma) supplemented with 10% Fetal Bovine Serum (FBS; Corning), 100 U/ml penicillin and 100 µg/ml streptomycin (1% Pen Strep; Sigma). Cells were incubated at 37 °C with 5% CO_2_.

### DNA constructs

All microtubule anchor constructs and the SspB-mCherry-p60 construct were cloned into a pB80 backbone containing 6 x GGGGS linkers in between protein segments described previously ^22^ using Gibson Assembly. Mouse katanin p60 described previously ^20^ was used for the SspB-mCherry-p60 construct. Human MAP7 and MAP7D3 coding regions described previously ^33^ were used as a template for iLID-mClover3-MAP7, iLID-mClover3-MAP7_N_(1-301), iLID-mClover3-MAP7_MTBD_(59-170), iLID-mClover3-MAP7D3, iLID-mClover3-MAP7D3_N_(1-309) and iLID-mClover3-MAP7D3_MTBD_(55-174). Human EB3 construct ^56^ was used as a template for EB3_N_-VVD-mClover3-iLID and EB3_N_-VVD-Halo-iLID constructs. Human 0N4R and 0N3R Tau construct ^57^ was used as a template for iLID-mClover3-Tau0N4R, iLID-mClover3-miniTau0N4R(140-342), iLID-mClover3-Tau0N4R MTBD(186-310) and iLID-mClover3-Tau0N3R. Rat MAP2C construct ^58^ was used as a template for iLID-mClover3-MAP2C and iLID-mClover3-MAP2C_MTBD_(300-393). Mouse DCX construct ^59^ was used as a template for iLID-mClover3-DCX. Rat CLIP115 from ^60^ was used as a template for CLIP115_N_(1-309)-mClover3-iLID. LAMP1-Halo was cloned by replacing mGFP with HaloTag in LAMP1-mGFP ^61^ using Gibson Assembly. Halo-Rab6A was cloned by replacing eGFP in eGFP-Rab6A ^62^ with HaloTag using Gibson Assembly. Halo-KDEL was cloned using Halo-Rab6A; A KDEL-encoding sequence (5’-AAAGACGAGCTATGA-3’) was inserted replacing Rab6A using overlapping primers and restriction enzyme cloning, and an ER targeting sequence derived from Heat Shock Protein family A member 5 (HSPA5; 5’-ATGAAGCTCTCCCTGGTGGCCGCGATGCTGCTGCTGCTCAGCGCGGCGCGGGCC-3’) was inserted directly in front of the HaloTag using Gibson Assembly.

### Cell transfection

Transfection of HeLa, COS-7 and U2OS was performed using FuGENE 6 (Promega) according to manufacturer’s instructions using a ratio of 3 µl FuGENE 6 per 1 µg DNA.

### Animals

Experiments were conducted in line with institutional guidelines of Utrecht University, and conducted in agreement with Dutch law (Wet op de Dierproeven, 1996) and European regulations (Directive 2010/63/EU). The animal protocol has been evaluated and approved by the national CCD authority (license AVD1080020173404). Female pregnant Wistar rats were obtained from Janvier, and embryos (both genders) at embryonic (E)18 stage of development were used for primary cultures of hippocampal neurons. The pregnant female rats were at least 10 weeks old and were not involved in any other experiments.

### Neuron isolation and culture

Rat primary hippocampal neurons were isolated from the hippocampi of embryonic day 18 pups as described previously ^63^ and plated on poly-L-lysine (Sigma) and laminin (Roche) coated coverslips. Primary neurons were cultured in Neurobasal medium (NB) supplemented with 2% B27 (Gibco), 0.5 mM glutamine (Gibco), 15.6 µM glutamate (Sigma), 100 U/ml penicillin and 100 µg/ml streptomycin (1% Pen Strep; Gibco). Cells were incubated at 37 °C with 5% CO_2_.

### Neuron transfection

Neurons were transfected on the 7^th^ day *in vitro* (DIV7) with 1 µg EB3_N_-VVD-mClover3-iLID, 2 µg SspB-mCh-p60 and 100 ng LAMP1-HaloTag using 3.3 µl Lipofectamine 2000 (Invitrogen), as described previously ^64^.

### HaloTag labelling

HaloTag dye JF646 (Promega) was diluted in DMSO to 200 µM, aliquoted and stored at −20 °C. Just prior to labelling a HaloTag aliquot was thawed and diluted 1:1000 in prewarmed cell media, vortexed for 5 s and incubated on cells for 15 min. Cells were washed 3 times with prewarmed cell culture PBS and then returned to cell media, live cell imaging was conducted over the next 5 hours.

### Microscopy

#### Spinning Disc

Spinning disk confocal microscopy was performed on an inverted research microscope Nikon Eclipse Ti-E (Nikon), equipped with the perfect focus system (Nikon), Nikon Plan Apo VC 100x N.A. 1.40 oil objective (Nikon) and a spinning disk-based confocal scanner unit (CSU-X1-A1, Yokogawa). The system was also equipped with ASI motorized stage with the piezo plate MS-2000-XYZ (ASI), Photometrics PRIME BSI back illuminated sCMOS camera (version USB 3, Teledyne Photometrics) and controlled by the MetaMorph 7.10 software (Molecular Devices). Lasers were used as the light sources: 488 nm 150 mW (Vortran Stradus 488, Vortran Laser Technology), 561nm 100 mW (OBIS 561-100LS, Coherent) and 639 nm 150 mW (Vortran Stradus 639, Vortran Laser Technology). We used ET-GFP filter set (49002, Chroma) for imaging of proteins tagged with mClover3; ET-mCherry filter set (49008, Chroma) for imaging of proteins tagged with mCherry; ET-Cy5 filter set (49006, Chroma) for imaging of proteins tagged with HaloTag JF646. 16-bit images were projected onto the sCMOS camera chip at a magnification of 63 nm/pixel. To keep cells at 37°C we used stage top incubator (model INUBG2E-ZILCS, Tokai Hit). For localized illumination iLas system was used and controlled with iLas software (Gataca Systems).

#### TIRF

Azimuthal TIRFM was performed on an inverted research microscope Nikon Eclipse Ti-E (Nikon), equipped with the perfect focus system (Nikon), Nikon Apo TIRF 100x N.A. 1.49 oil objective (Nikon) and iLas3 system (Dual Laser illuminator for azimuthal spinning TIRF (or Oblique) illumination and Simultaneous Targeted Laser Action including PhotoAblation; Gataca Systems). The system was also equipped with ASI motorized stage MS-2000-XY (ASI), Photometrics CoolSNAP Myo CCD camera (Teledyne Photometrics) and controlled by the MetaMorph 7.8 software (Molecular Devices). Stradus 488 nm (150 mW, Vortran), OBIS 561 nm (100 mW, Coherent) and Stradus 642 (110 mW, Vortran) lasers were used as the light sources. We used ZT405/488/561/640rpc ZET405/488/561/635m filter set (TRF89901, Chroma) together with Lambda 10-3 (Sutter Instrument) filter wheel, equipped with the emission filters ET630/75m and ET700/75m (from ET-mCherry 49008 and ET-Cy5 49006 filter sets, Chroma). 16-bit images were projected onto the CCD chip at a magnification of 45.4 nm/pixel. To keep cells at 37°C we used stage top incubator (model INUBG2E-ZILCS, Tokai Hit). Localized illumination was controlled with iLas software (Gataca Systems).

#### Widefield Microscopy

Widefield imaging was performed using a Nikon Eclipse 80i upright microscope fitted with a Photometrics CoolSNAP HQ2 CCD camera and a CoolLED illumination system. Microscope was fitted with a set of Chroma filters, the following filters were used to image samples: ET-BFP2 (49021) to image Alexa Fluor Plus 405, ET-GFP (49002) to image mClover3, ET-mCherry (49008) to image mCherry, ET-Cy5 (49006) to image Alexa Fluor 647. A Plan Apo VC 60x N.A. 1.40 oil objective was used to image samples and Nikon NIS Br software was used to control the microscope.

#### Fixed Confocal Microscopy

For fixed confocal imaging a Carl Zeiss LSM880 Fast AiryScan microscope fitted with 405nm, Argon Multiline, 561nm and 633 nm lasers and AiryScan and PMT detectors was used. A Alpha Plan-APO 100x/1.46 Oil DIC VIS objective was used to image samples and ZEN 2.3 software was used to control the microscope.

#### Live cell opto-katanin microtubule severing

U2OS/COS-7/HeLa cells were seeded at 15% confluency on a 25 mm coverglass inside one well of a 6 well plate one day prior to transfection. Cells were transfected with 1 µg EB3_N_-VVD-mClover3-iLID and 2 µg SspB-mCh-p60 and co-transfected with 50 ng HaloTag-Rab6A or 100 ng HaloTag-KDEL or incubated with 100 nM SiR tubulin in cell media overnight, then used for the experiment the following day, taking care to protect cells from blue, violet and UV light after transfection. Cells were labelled with HaloTag dye if applicable, then imaged on a Spinning Disc microscope (or TIRF microscope in the case of ER imaging), a 561 nm laser was used to image SspB-mCh-p60 with one 500 ms exposure (0.41 mW, 500 µW/cm^2^) every 4 s or 60 s, a 642 nm laser was used to image JF646 HaloTag dyes or SiR tubulin with one 500 ms exposure (0.25 mW, 300 µW/cm^2^) every 2 or 4 s, for localized activation a FRAP unit was used to apply one localized 113 ms pulse of 488 nm light (3 µW, 700 µW/cm^2^) every 4 s. In the case of mClover3 anchor imaging and field of view activation, a 488 nm laser was used with one 500 ms exposure (0.25 mW, 300 µW/cm^2^) every 4 s. Quantification of local fluorescence intensity was performed in Fiji, quantification of ER nodules was done manually in MetaMorph 7.10.2.240, data was normalized in Excel, data plots were prepared in Graphpad Prism.

#### Immunofluorescence

For fixed cell analysis of microtubule anchors, HeLa cells plated on coverslips transfected the day prior were either fixed in the dark (DARK), or first placed in an incubator under blue LED lights (800 µW/cm^2^) for 16 minutes prior to fixation (LIT). Cells were first fixed with Methanol at −20°C for 7 minutes, followed by fixation with 4% paraformaldehyde in PBS for 15 minutes at room temperature. Cells were permeabilized with 0.2% Triton-X in PBS for 2.5 minutes. Neurons were instead fixed with 4% paraformaldehyde with 1% sucrose in MRB80 buffer prewarmed to 37°C for 12 minutes at room temperature. Neurons were subsequently washed 3 times with PBS, then permeabilized with 0.1% Triton-X in PBS for 10 minutes at room temperature. Samples were blocked for 1 hour in blocking buffer consisting of 2% Bovine Serum Albumin with 0.05% Tween-20 in PBS before application of primary antibodies. Primary antibodies were diluted 1:200 in blocking buffer and incubated on samples for 1 hour, antibodies used were: Mouse anti-α-tubulin (Sigma, T6199), Rat anti-tyrosinated-α-tubulin (Thermo, MA1-80017), Mouse anti-acetylated-tubulin (Sigma, T7451). Samples were washed 3 times with PBS then incubated with highly cross-preadsorbed secondary antibodies diluted 1:250 in blocking buffer and incubated for 1 hour, secondaries used were goat anti-mouse conjugated with Alexa Fluor Plus 405 (Thermo Fisher, A48255) and goat anti-rat conjugated with Alexa Fluor 647 (Thermo Fisher, A21247). Samples were washed 3 times with PBS and mounted with Vectashield (Vector Labs). HeLa cells transfected with different anchors and stained for α-tubulin were imaged with a widefield Nikon upright microscope. COS-7 cells and neurons stained for different microtubule populations or α-tubulin were imaged on a Zeiss LSM880 Fast AiryScan confocal microscope.

## Data analysis

### Fluorescence intensity normalization

For live cell imaging experiments a neighboring transfected or SiR-tubulin labelled cell was used for normalization to account for photobleaching over time. In the case of fixed and stained samples tubulin staining and mCherry/mClover3 intensity was normalized to untransfected neighboring cells. This was done using the following equation: 

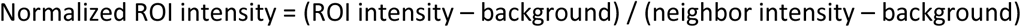

For live cell fluorescence intensity over time plots, the normalized fluorescence intensities were subsequently divided by the average of the first 5 measurements to also account for variability in construct expression or SiR-tubulin staining.

### Dynamic lysosome analysis

Non-dynamic lysosomes were first filtered out using the Faster Temporal Mean plugin (https://github.com/marcelocordeiro/MedianFilter-ImageJ). Dynamic lysosomes were then counted inside the ROI where microtubules were severed and in a comparable region where microtubules were not severed, using the ComDet plugin (https://github.com/ekatrukha/ComDet). Dynamic lysosome counts were normalized to the average of the first 10 frames.

### Membrane dynamics analysis

Golgi and ER membrane dynamics were quantified using the Membrane Displacement Analysis Fiji Macro developed and described previously ^65^, kindly shared with us by L. M. Voortman (Leiden University Medical Centre, Leiden, The Netherlands). The output of this optical flow algorithm expresses membrane displacement in pixel intensity where a higher intensity corresponds to a higher displacement and an intensity of 0 corresponds to no displacement. Therefore in order to express membrane dynamics within a ROI, the mean pixel intensity was quantified and normalized to the average of the first 10 time points.

### Statistical analysis

A Mann-Whitney U test was used to test for statistical significance between DARK and LIT treatment groups. A P value below 0.05 was regarded as statistically significant.

### Data availability

All data that support the conclusions are included in the manuscript or available from the authors on request.

## Acknowledgements

The authors would like to acknowledge: L. M. Voortman for sharing and trouble-shooting use of the Membrane Displacement Analysis macro; P. J. Hooikaas for advising on MAP7 constructs; R. J. McKenney for advising on Tau constructs; E. A. Katrukha for advising on lysosome dynamics analysis; M. Siemons for advising on laser power calculations. J.C.M.M was supported by an EMBO Long Term fellowship number ALTF 261-2019 and A.A. was supported by ZonMW Top grant 91216006.

## Author Contributions

J.C.M.M. wrote the manuscript, cloned constructs, performed or coordinated experiments and analysis and acquired funding, I.G. performed fixed anchor construct imaging and analysis and conducted ER TIRF experiments and analysis, W.N. advised on optogenetic modules and shared reagents, L.C.K. acquired funding and supervised, A.A. supervised the study, reviewed the manuscript and acquired funding.

## Extended Data

**Extended Data Fig. S1:**
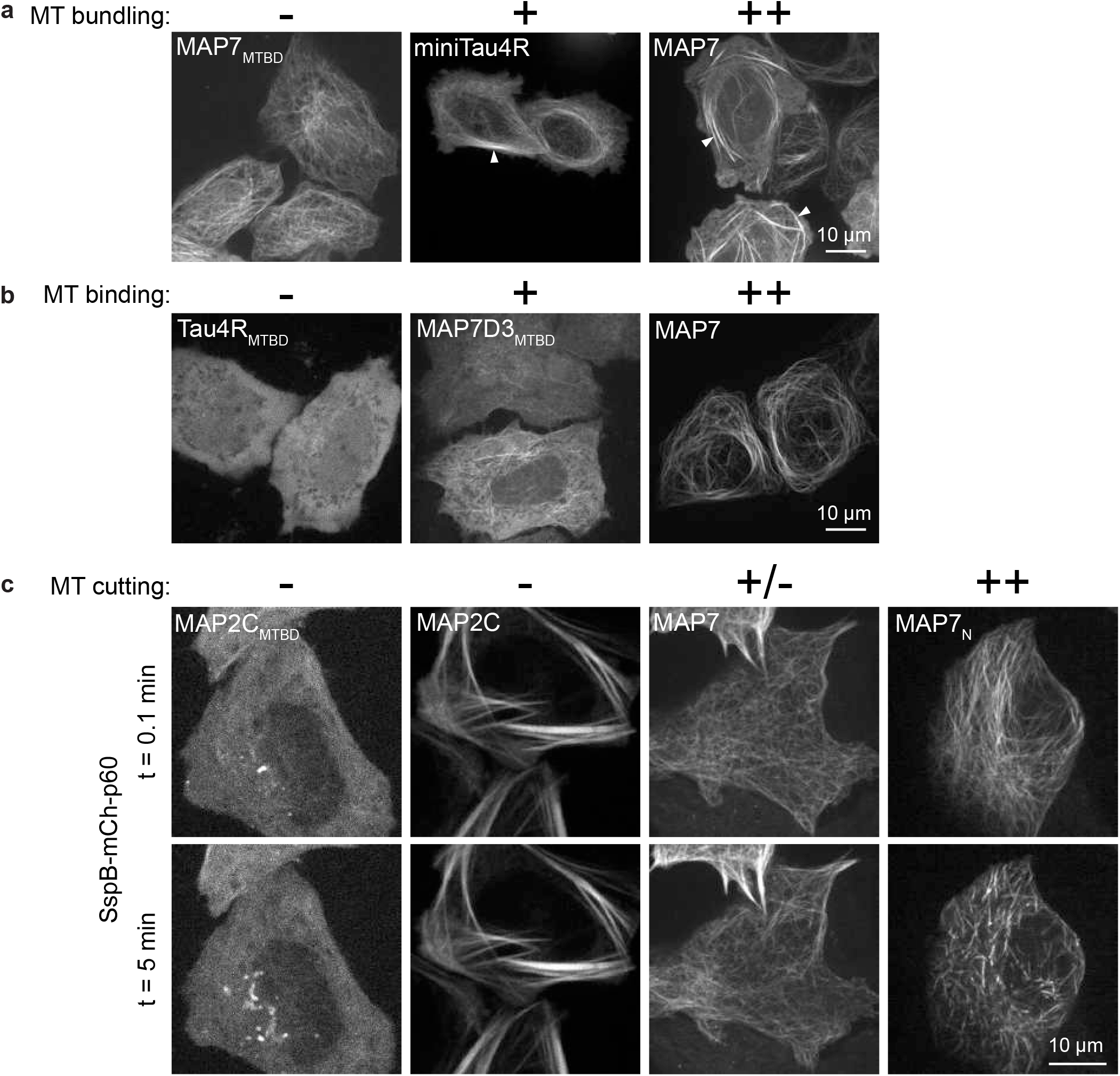
Classification of anchor constructs according to microtubule binding, bundling and severing with SspB-mCh-p60. **a**-**c**, Anchor constructs were co-transfected in HeLa cells along with SspB-mCh-p60. Live cells were imaged for 5 minutes to visualize both SspB-mCh-60 (561 nm) and mClover3 labelling the anchor (488 nm), thus activating iLID-SspB, on a spinning disc microscope. **a,b**, Based on the localization of the anchor in the first frame the (**a**) microtubule bundling (indicated by arrows) and (**b**) microtubule binding properties of the anchors were assessed to be either high (++), moderate (+) or not observed (-), one example is shown for each category. **c**, Based on microtubule severing observed in the mCherry channel over the course of 5 minutes, the potential of the anchor to sever microtubules by recruitment of SspB-mCh-p60 was judged to be high (++), poor (+/-) or not observed (-). An example is shown for each category. In cases where no severing was observed, this could be due to poor microtubule binding of the anchor and poor recruitment of SspB-mCh-p60 (MAP2C_MTBD_); however, some anchors could bind microtubules and recruit SspB-mCh-p60 but did not enable microtubule severing (MAP2C). See Table 1 for results.

**Extended Data Fig. S2:**
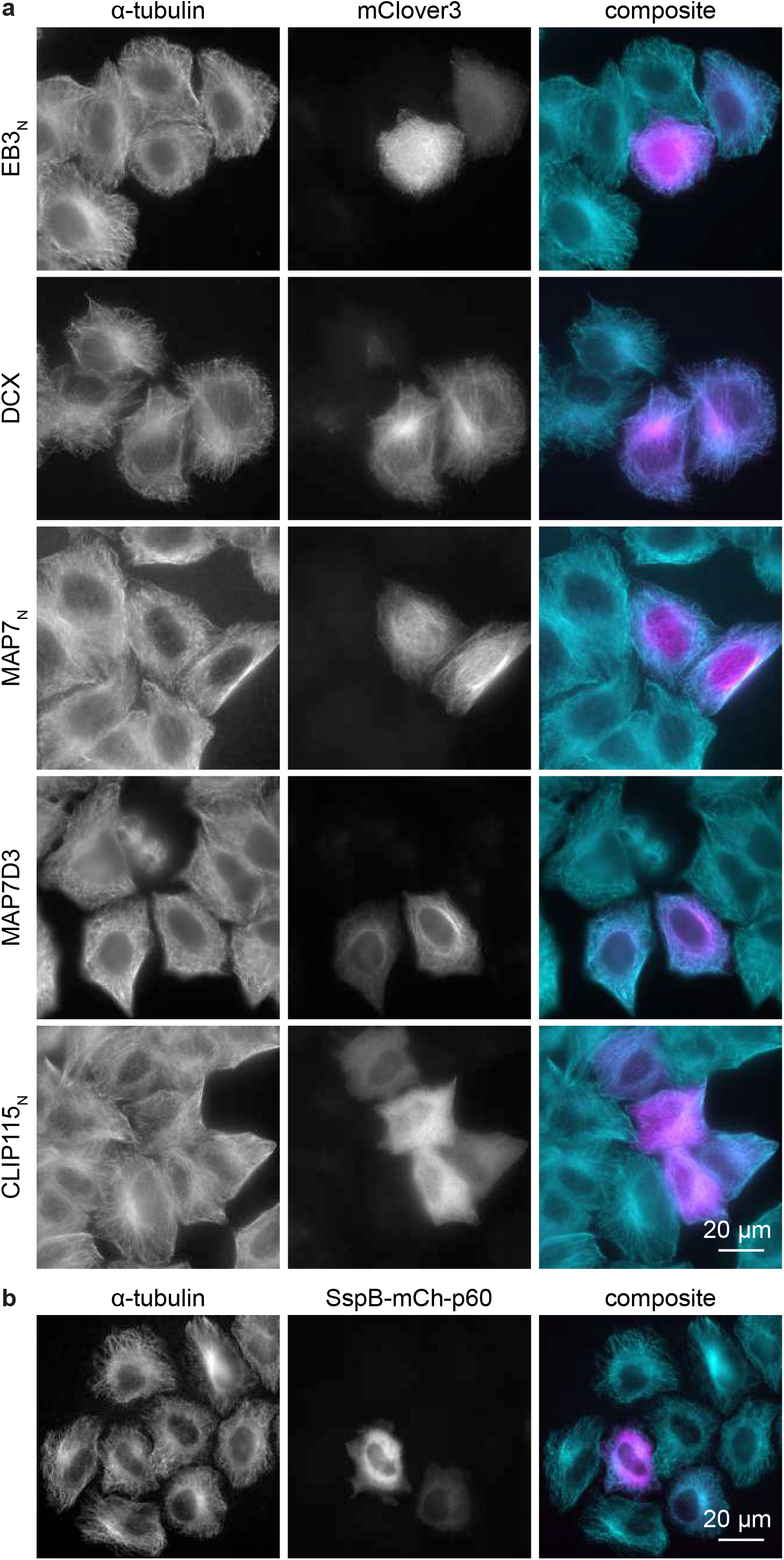
Characterization of the effect of anchor constructs or SspB-mCh-p60 on microtubule organization and staining. **a,b**, HeLa cells transfected with (**a**) anchor constructs or (**b**) SspB-mCh-p60 stained for α-tubulin.

## Supplementary Videos

**Supplementary Video 1: Localized co-recruitment of optogenetic constructs**. U2OS cell co-expressing SspB-mCh-p60 (left) and EB3_N_-VVD-mClover3-iLID (right) locally pulsed with 488 nm in the area outlined by a blue square for a period of 1.5 min, indicated by a blue dot in the top right corner.

**Supplementary Video 2: Microtubule severing around the centrosome and near the cell periphery**. U2OS cell co-expressing SspB-mCh-p60 (magenta) and EB3_N_-VVD-mClover3-iLID (not visualized), labelled with SiR-tubulin (cyan) locally pulsed with 488 nm in the areas outlined by yellow squares from 8 s onward, as indicated by a blue dot in the top right corner.

**Supplementary Video 3: Microtubule severing followed by microtubule recovery**. U2OS cell co-expressing SspB-mCh-p60 (left) and EB3_N_-VVD-mClover3-iLID (not visualized), labelled with SiR-tubulin (right) locally pulsed with 488 nm in the area outlined by a blue square for a period of 1.5 min, indicated by a blue dot in the top right corner.

**Supplementary Video 4: Localized microtubule severing in neurites**. DIV9 neuron co-expressing SspB-mCh-p60 (left) and EB3_N_-VVD-mClover3-iLID (not visualized), labelled with SiR-tubulin (right) locally pulsed with 488 nm in the areas outlined by the blue boxes from 20 s onward, as indicated by a blue dot in the top right corner.

**Supplementary Video 5: Blocking endolysosome transport in neurons by localized microtubule severing**. DIV9 neuron co-expressing SspB-mCh-p60, EB3_N_-VVD-mClover3-iLID and late endosome and lysosome marker LAMP1-HaloTag (shown). Cell was locally pulsed with 488 nm in the area outlined by the blue box from 10 min onward, as indicated by a blue dot in the top right corner.

**Supplementary Video 6: Blocking exocytic Rab6A vesicle transport by localized microtubule severing**. U2OS cell co-expressing SspB-mCh-p60 (magenta), EB3_N_-VVD-mClover3-iLID (not visualized) and exocytic vesicle and Golgi marker Halo-Rab6A (cyan). Cell was locally pulsed with 488 nm in the area outlined by the yellow box from 2 min onward, as indicated by a blue dot in the top right corner. White box indicates zoomed area shown in the top center. White arrows highlight examples of pausing vesicles.

**Supplementary Video 7: Local perturbation of ER tubulation by targeted microtubule severing**. COS-7 cell co-expressing SspB-mCh-p60 (magenta), EB3_N_-VVD-mClover3-iLID (not visualized) and ER marker KDEL-HaloTag (cyan). Cell was locally pulsed with 488 nm from 5 min onward, as indicated by a blue dot in the top right corner, resulting in microtubule disassembly within the area outlined by the yellow circle.

**Supplementary Video 8: Local perturbation of Golgi stack and Rab6A vesicle dynamics by targeted microtubule severing**. U2OS cell co-expressing SspB-mCh-p60 (magenta), EB3_N_-VVD-mClover3-iLID (not visualized) and exocytic vesicle and Golgi marker Halo-Rab6A (cyan). Cell was locally pulsed with 488 nm from 10 min onward, as indicated by a blue dot in the top right corner, resulting in microtubule disassembly within the area outlined by the yellow circle. Arrows highlight Golgi tubules, absent after microtubule severing.

**Supplementary Video 9: Relaxation of nuclear deformation by sequential severing of constricting microtubule bundles**. COS-7 cell co-expressing SspB-mCh-p60 (shown) and EB3_N_-VVD-mClover3-iLID (not visualized). Cell was locally pulsed with 488 nm within the regions indicated by the yellow boxes, pulsing events are occurring when boxes are shown in blue.

